# Cortico-motor control dynamics orchestrates visual sampling

**DOI:** 10.1101/2020.03.23.003228

**Authors:** Alice Tomassini, Eric Maris, Pauline Hilt, Luciano Fadiga, Alessandro D’Ausilio

## Abstract

Movements overtly sample sensory information, making sensory analysis an active-sensing process. In this study, we show that visual information sampling is not just locked to the (overt) movement dynamics, but it is structured by the internal (covert) dynamics of cortico-motor control. We asked human participants to perform an isometric motor task – based on proprioceptive feedback – while detecting unrelated near-threshold visual stimuli. The motor output (Force) shows zero-lag coherence with brain activity (recorded via electroencephalography) in the beta-band, as previously reported. In contrast, cortical rhythms in the alpha-band systematically forerun the motor output by 200ms. Importantly, visual detection is facilitated when cortico-motor alpha (not beta) synchronization is enhanced immediately before stimulus onset, namely at the optimal phase relationship for sensorimotor communication. These findings demonstrate an automatic gating of visual inputs by the ongoing motor control processes, providing evidence of an internal and alpha-cycling visuomotor loop.

## Introduction

Rather than being serially ordered along distinct processing stages, action and perception are now deemed to be tightly intertwined along all of the processing stages (1). The latter take the form of loops, whereby descending (motor) signals interact at multiple timescales with different internal (predictions) and ascending (sensory) signals that inform about the body (e.g. proprioception) and the external world (e.g. vision). Effective behavior indeed relies on a dynamic interplay between multimodal sensorimotor loops (2). To close the sensorimotor loop, the sampling of sensory inputs must be finely synchronized to the issuing of descending commands/predictions.

Oscillatory activity is likely to coordinate the information flow across sensorimotor circuits (3–5). Neuronal excitability is subject to ongoing oscillatory fluctuations which yield measurable behavioral consequences, at both perceptual (6–9) and motor (10, 11) levels. Phase synchronization of these oscillatory dynamics further provides a mechanism to appropriately time the excitability of distant groups of neurons, enabling functional coupling and selective information exchange (12–14).

Recent evidence suggests that oscillations may synchronize action onset and perceptual sensitivity. Cortical excitability (15–20) and visual performance (18, 20–25) undergo rhythmic modulations that are time-locked to movement onset (see 26 for a review). Such a perceptual/neural modulation begins before actual motor production and affects stimulus processing even when unrelated to the motor task. This phenomenon is not, however, necessarily *motor* in origin. The probability of initiating a movement could be itself modulated by a spontaneous, sensory-driven or top-down controlled neural oscillatory dynamics that is not otherwise involved in motor computations, i.e., in the preparation and control of movements. Indeed, some accounts posit that perceptual sensitivity and movement initiation are jointly coordinated in time by an attention-based clocking mechanism (23, 27–29). At present, there is no compelling evidence that the *internal* motor control dynamics bears direct relevance for (visual) sensory analysis.

Here we put this hypothesis to a test. To this aim, we turned to a task requiring continuous (proprioceptive-based) closed-loop control of the motor output (isometric force control) and examined the ongoing relationship between the motor output (Force) and brain activity recorded via electroencephalography (EEG). This relationship – classically described in the spectral domain as corticospinal coherence (30, 31) – provides a privileged window into the dynamics of cortico-motor control. We thus tested whether visual detection of stimuli unrelated to the motor task is inherently coupled to the ongoing dynamics of cortico-motor control, as indexed by cortico-force coherence.

## Results

We recorded EEG, EMG and force on twenty healthy human participants who were asked to perform two tasks concurrently: continuous isometric contraction and visual detection. Participants applied force with their right hand on an isometric joystick until they reached the required force level with the aid of visual feedback (see Methods for details); henceforth, they were instructed to keep tonic contraction for 5.5 s without feedback while waiting for the appearance of a visual dot with near-threshold contrast. The visual stimulus appeared at a random time (ranging from 1.6 to 4.6 s) in 85% of trials. At the end of the trial, a question mark appeared on the screen which signaled participants to release the force and report verbally whether they had seen or not seen the stimulus (Fig 1A,B).

**Fig 1.**
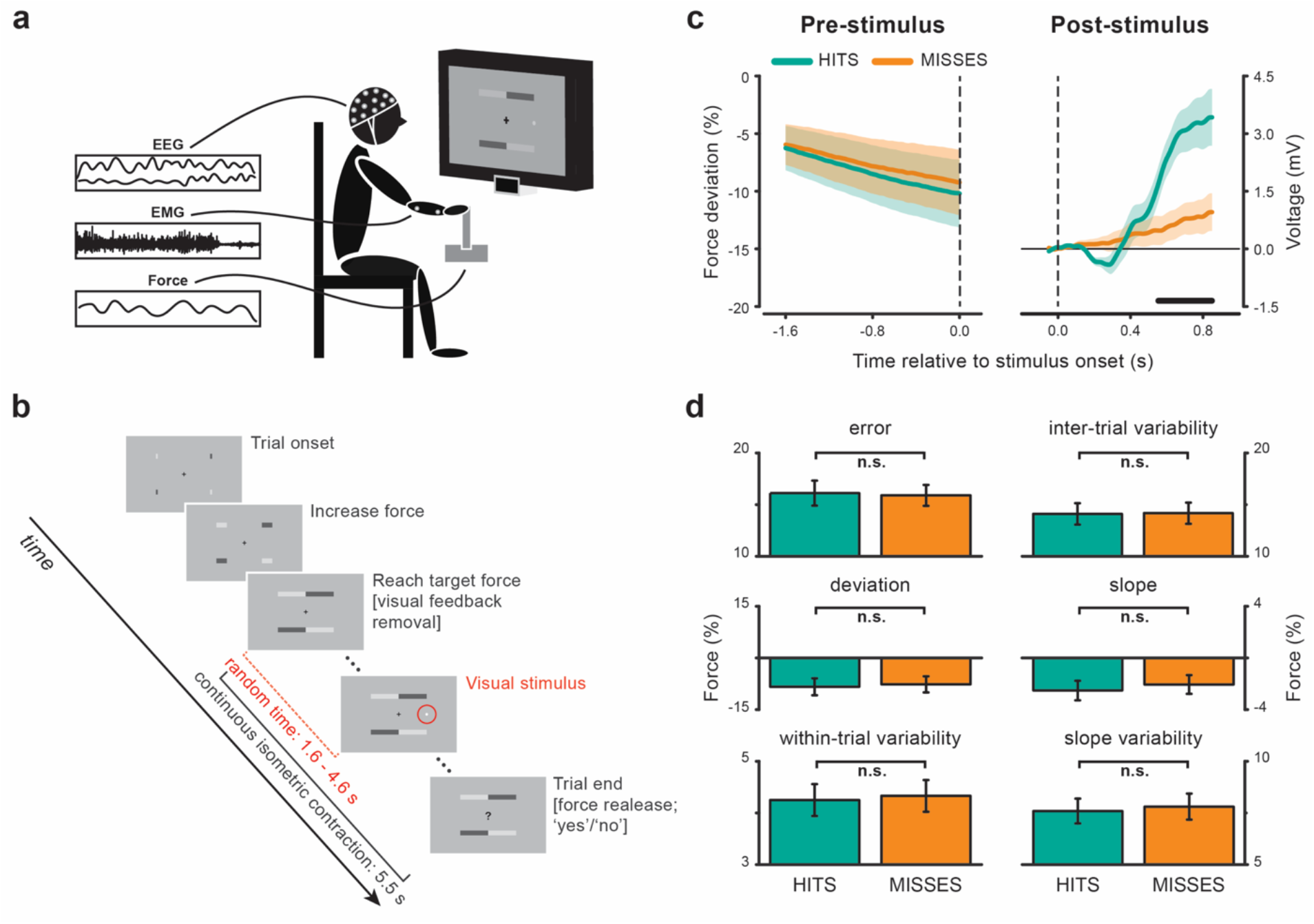
Experimental set-up, procedure and behavioral results. (**A**) EEG, EMG and Force were recorded while participants performed two tasks concurrently: visual detection and right wrist abduction to push an isometric joystick’s handle towards one’s own body. (**B**) Visual feedback of the force (four horizontal bars elongating towards the center of the screen) was provided until participants reached the target force level (see Methods). Afterwards, participants were required to maintain stable contraction for 5.5 s without visual feedback. During continuous contraction, a near-threshold visual dot could appear 7.5° to the right of fixation and at a random time between 1.6 and 4.6 s (no stimulus was presented in 15% of trials). Trial end was signaled by a question mark prompting participants to release the contraction and report verbally whether they had seen or not seen the visual stimulus. (**C**) Force time courses in the pre- (left) and post- (right) stimulus period for hits- and misses-trials. Shaded areas represent ± 1 standard error of the mean (SEM). The black horizontal line indicates the time interval (0.55-0.85 s) belonging to the cluster that survived cluster-based permutation statistics for the hits-misses contrast. (**D**) Motor performance in the −1.6-0-s-window before stimulus presentation quantified as absolute (error) and relative (deviation) percentage difference from target force, intertrial and within-trial force variability, slope and slope variability. Error bars represent ± 1 SEM.

### Behavioral performance

By design, performance in the visual task is at threshold (hits – ‘yes’ responses for stimulus-present trials: 47±5%; misses – ‘no’ responses for stimulus-present trials: 52±5%; MEAN±SD) and shows a low rate of false alarms (‘yes’ responses for stimulus-absent trials: 1.3±1.7%). To investigate whether performance in the perceptual and motor task co-varied, we analyzed the force time course separately for hits and misses. Fig 1C shows the time course of the force in the pre-(left) and post-stimulus (right) epochs. Overall, pre-stimulus force is ~8% lower than the instructed force level (t_19_ = −3.3565, p = 0.0033) and shows a slowly declining temporal trend (mean slope: −2.27%; t_19_ = −3.1017, p = 0.0059). However, produced force is not at any time point different between hits and misses (Fig 1C, left). Many metrics describing force level and variability [including absolute (error; p = 0.5061) and relative (deviation; p = 0.1345) difference from target force, slope (p = 0.0547) and its variability (p = 0.1432), as well as intertrial (p = 0.8271) and within-trial force variability (p = 0.1570)] are comparable for hits- and misses-trials (Fig 1D). Trial-by-trial fluctuations in arousal and task engagement are expected to yield a positive correlation between perceptual and motor performance (e.g. higher arousal associated with performance improvements in both tasks); within-trial divided attention would instead produce a negative correlation (e.g. attention diverted to the motor task associated with better motor and worse perceptual performance). In contrast, force level and performance in the motor task do not show any type of co-variation with visual performance, suggesting that spontaneous fluctuations in attention/arousal had no major impact on behavior.

Unlike the pre-stimulus epoch, force is modulated by stimulus presentation in a detection-dependent fashion (Fig 1C, right). Specifically, the visual stimulus evokes a bi-phasic response with an initial negative deflection (from ~0.2 to ~0.4 s) which is visible only in hits-trials, followed by a later increase in force (from ~0.4 s onwards) which is significantly larger for hits compared to misses (p = 0.0174; permutation test corrected for multiple comparisons across time; cluster time interval: 0.55 – 0.85 s). These responses resemble the event-related potentials in the force shown previously by Novembre and colleagues (32) for supra-threshold somatosensory and auditory stimuli and will not be investigated further in the current study. The main analyses reported below were performed on the epoch preceding stimulus presentation time.

### Pre-stimulus cortico-force coherence

Our main aim was to investigate whether visual processing is coupled to the internal motor dynamics. To provide a window into the internal motor control processes, we investigated the relationship between the produced force (the outcome of the motor control processes) and the neural activity as measured using the EEG.

The force produced during continuous isometric contraction shows a distinctive rhythmic signature centered at ~11 Hz (Fig 2A left), which is commonly designated as physiological tremor (33).

**Fig 2.**
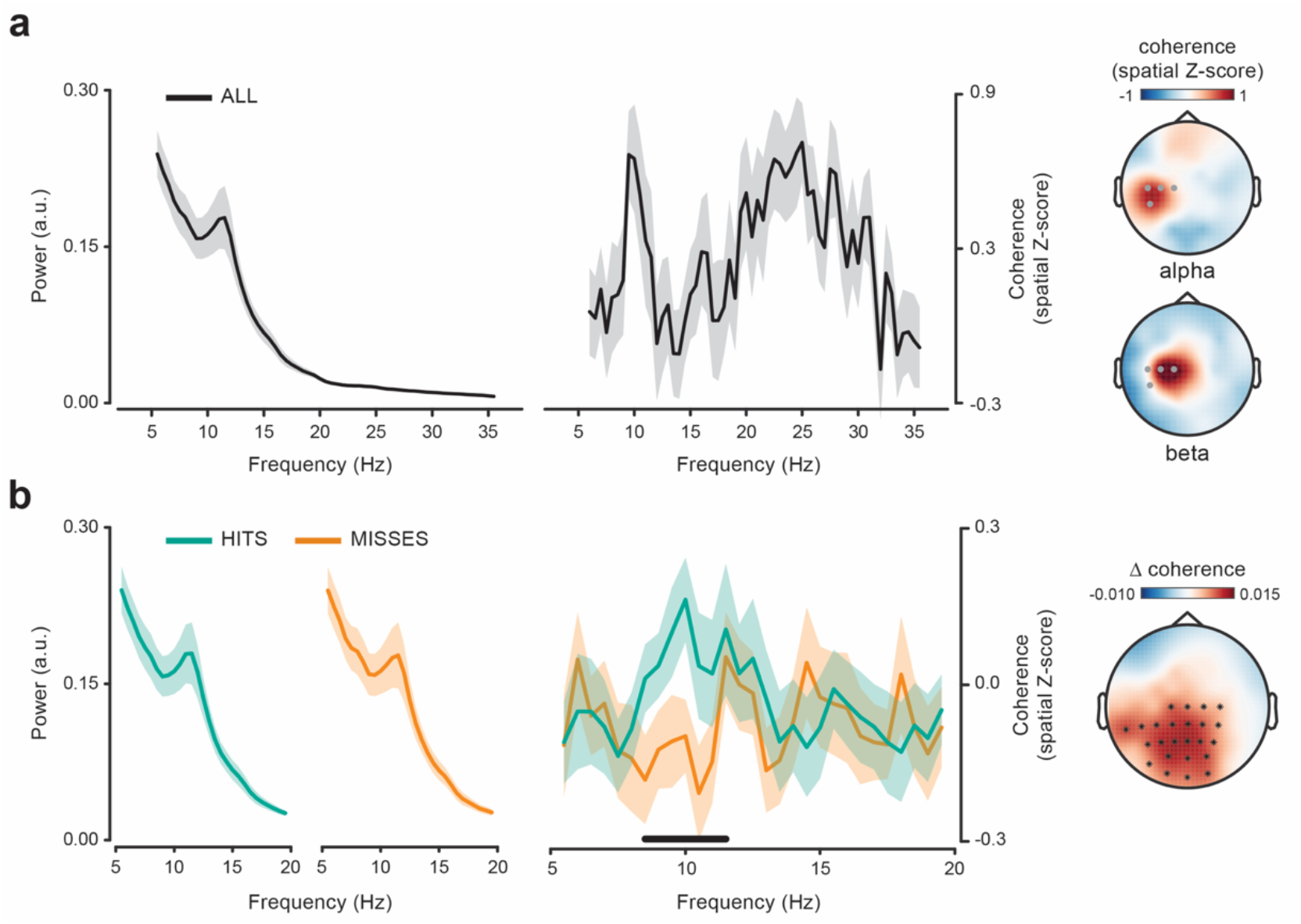
Spectral content of pre-stimulus force and coherence with cortical activity. (**A**) Force power (left) and coherence (middle) spectrum with contralateral centro-parietal EEG electrodes (C1, C3, C5, CP5; marked in grey in the topographic maps) computed on the pre-stimulus window (−1.6-0 s). Topographies show coherence in the alpha (9-11 Hz; top) and beta (20-30 Hz; bottom) range. Coherence has been spatially z-scored (34) before averaging across subjects and frequencies. (**B**) Same as in (A) but computed separately for hits- and misses-trials (left, middle). The black horizontal line indicates the frequency interval (8.51-1.5 Hz) belonging to the cluster that survived cluster-based permutation statistics for the hits-misses contrast. Coherence spectra are averaged over the EEG electrodes belonging to the same cluster (evaluated at 10.5 Hz; see black asterisks in the topographic map). Topography shows the hits-misses difference in coherence averaged over the cluster frequency interval (8.5-11.5 Hz).

First, we examined whether this peripheral rhythm in the force is related to a central rhythm by computing phase coherence (see Methods). Fig 2A (middle) shows that the cortico-force coherence spectrum has a clear peak around 10 Hz. Coherence in the alpha-band is spatially confined to the contralateral side and reaches maximal values at centro-parietal electrodes (CP5, C5; Fig 2A, topography on the top; coherence spatially z-scored and averaged over subjects and frequencies between 9 and 11 Hz).

Coherence between peripheral (especially muscular) and cortical activity has been most commonly reported in the beta-band (see 30, 35, 36 for reviews), and much less frequently in the alpha-band (37–39). Despite the force signal does not display a distinct beta-band spectral peak (Fig 2A, left), we do observe a consistent increase in cortico-force coherence also in this frequency range (40; Fig 2A, middle). Though partly overlapping, the topography for beta coherence is distinct from that for alpha coherence, as it peaks over more medial and anterior electrodes (C1, C3; Fig 2A, topography on the bottom; 20-30 Hz), in full agreement with what typically observed for cortico-muscular coherence (13, 41). This suggests that at least partially segregated neuronal populations are involved in alpha and beta coherence.

We next assessed whether hits and misses could be differentiated based on the pre-stimulus cortico-motor state as evaluated through cortico-force coherence. Alpha-band fluctuations in the force are more consistently synchronized with central rhythmic activity before hits compared to misses. Specifically, hits are preceded by stronger cortico-force coherence over a large array of central and posterior electrodes (cluster p = 0.005; cluster frequency interval: 8.5-11.5 Hz; corrected for frequencies between 5 and 35 Hz; Fig 2B). This modulation is exclusive to the alpha range as no difference is observed within the beta range or at any other tested frequency. Moreover, it cannot be explained by differences in alpha tremor amplitude which is comparable irrespective of the perceptual outcome (Fig 2B, left). The effectiveness of alpha phase synchronization within the motor system is thus relevant for visual perception (in the absence of changes in motor output; see above).

### Lagged cortico-force coherence

From a mechanistic point of view, one important aspect of the cortico-muscular/force coherence is the directionality of the interaction between cortical structures and motor output. A possible way to investigate this directionality is to look at the dependence of coherence on the relative timing (lag) between signals. We thus investigated cortico-force coherence by systematically varying the lag between the force and the EEG signal. We computed this lagged coherence between a fixed 0.6-s force segment (from −1.1 to −0.5 s relative to stimulus onset) and 0.6-s EEG segments that were time-shifted (relative to the force signal) by lags ranging from −0.5 s (EEG precedes force) to +0.5 s (EEG follows force) in steps of 10 ms. Fig 3A shows the lag-frequency representation of cortico-force coherence for two selected electrodes that better capture coherence either in the beta-(C1; left) or in the alpha- (CP5; right) band. While coherence in the beta-band is concentrated around lag zero, coherence in the alpha-band is clearly biased towards negative lags, reaching maximal values in the lag interval between – 0.2 and −0.15 s (see S1 Fig for similar results on cortico-EMG coherence).

**Fig 3.**
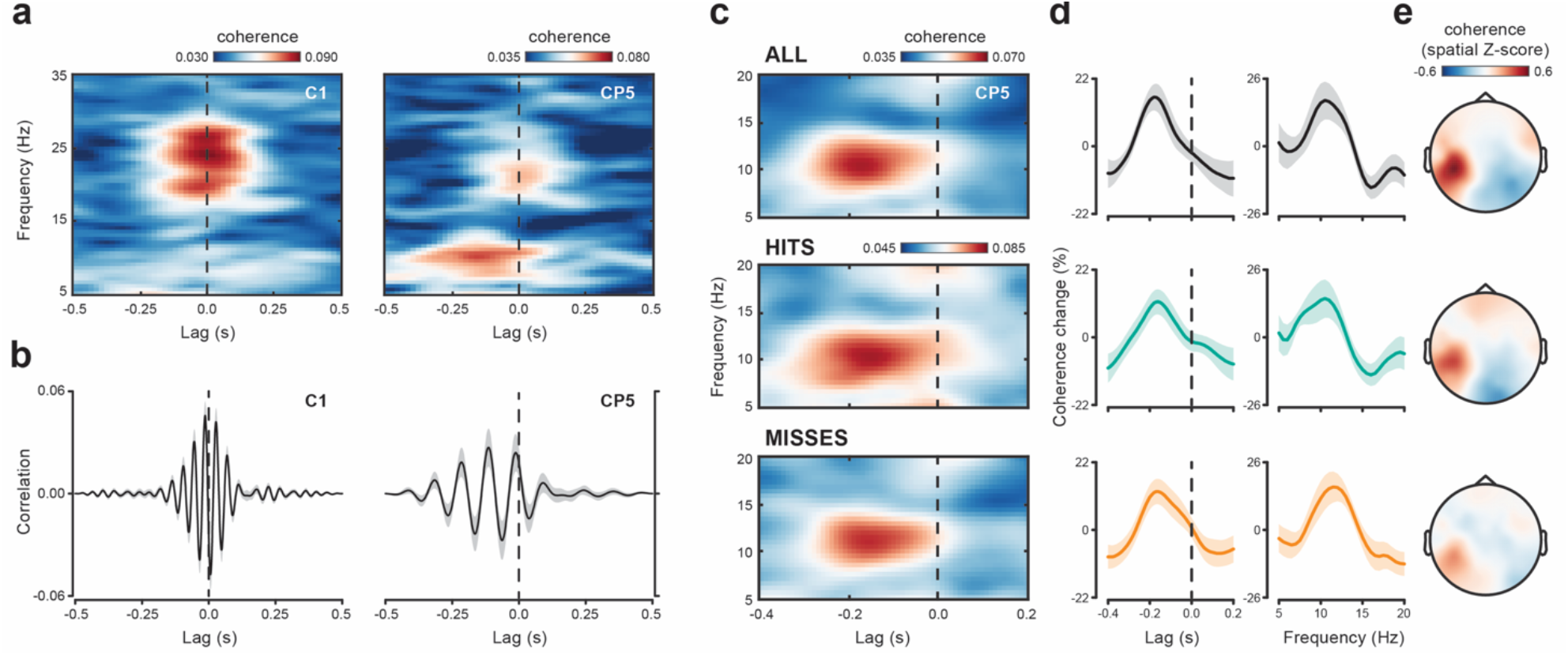
Lag-dependency of cortico-force coherence. (**A**) Lag- and frequency-resolved cortico-force coherence is shown for two EEG electrodes, C1 (left) and CP5 (right), where beta- and alpha-band coherence is maximal, respectively. Coherence has been calculated on 0.6-s data windows (from −1.1 to −0.5 s) by shifting the EEG signal (relative to the force signal) by a variable amount of time (negative lags: EEG precedes force; zero-lag: EEG and force are time-aligned; positive lags: EEG follows force) (**B**) Cross-correlations between force and EEG activity [over the same electrodes as in (A)] that were previously band-pass filtered in the beta (20–30 Hz; left) and alpha (8–12 Hz; right) range. Cross-correlations are normalized so that the autocorrelations at zero lag are identically 1. (**C**) Lag-frequency coherence representation as in (A) but computed on shorter (0.3-s) sliding data windows and then averaged over the pre-stimulus period for all trials as well as separately for hits- and misses-trials. (**D**) Lag (left) and spectral (right) tuning of cortico-force coherence expressed as the relative percentage change in coherence averaged over frequencies between 9 and 12 Hz and lags between −0.22 and −0.14 s, respectively. (**E**) Topographies show coherence at frequency 10.5 Hz and lag −0.2 s for all trials (top), hits (middle) and misses (bottom).

This peculiar lag-dependency profile is also visible in the cross-correlation functions between 1-s force and EEG segments (from −1 to 0 s) that were previously band-pass filtered in the relevant frequency ranges (beta-band: 20-30 Hz; alpha-band: 8-12 Hz; two-pass Butterworth filter, 2^nd^ order for each single pass). Fig 3B shows that correlation between the beta-band filtered signals is symmetrical around zero lag (left). Conversely, the correlation between the alpha-band filtered signals is clearly asymmetrical and leftward-shifted with respect to lag zero (right). These results indicate that the EEG alpha rhythm foreruns a corresponding peripheral rhythm in the force by about 0.2 s.

To increase the temporal resolution of our analyses we computed again lagged coherence but now using a shorter 0.3-s sliding time window that was advanced over the pre-stimulus epoch (from −1 to −0.35 s) in 10-ms steps. Moreover, to evaluate the consistency in the properties of alpha-band coherence across different trial categories, we performed the same analysis for all trials as well as separately for hits and misses.

Fig 3C shows the results of this analysis collapsed over the time dimension (i.e., averaged over the pre-stimulus epoch), restricted to the frequency range between 5 and 20 Hz and to the lag range between −0.4 and +0.2 s. Coherence (evaluated at electrode CP5) shows selective spectro-temporal properties, being higher in the alpha-band than in the other frequencies, and higher at negative than at positive lags. In particular, the lag-dependency profile peaks at −0.18 s for all trials and at a very similar lag for hits and misses (−0.16 s) (Fig 3D, left; relative % change in coherence averaged for frequencies between 9 and 12 Hz). The spectral profile is also comparable across trials subsets with maximal coherence values observed at 10.5 Hz for all trials, 10.5 Hz for hits and 11.5 Hz for misses (Fig 3D, right; relative % change in coherence averaged for lags between −0.22 and −0.14 s). Finally, the scalp topography of alpha-band coherence is very similar across hits and misses and closely resembles the topography already obtained by computing (non-lagged) coherence over a much longer time window (> 1 s; see Fig 2A).

### Time-dependent changes in the perceptual relevance of alpha coherence

We next applied the time-resolved approach to investigate whether the contribution of alpha-band coherence to perceptual performance varies over the pre-stimulus window. Indeed, if the ongoing state of cortico-force coupling is relevant for perceptual processes we could expect its impact to be maximal as we get closest to the stimulus presentation time.

Fig 4 shows the time- and lag-resolved coherence calculated for both hits and misses at 10.5 Hz (i.e., the frequency of maximal coherence for all trials combined; see above).

**Fig 4.**
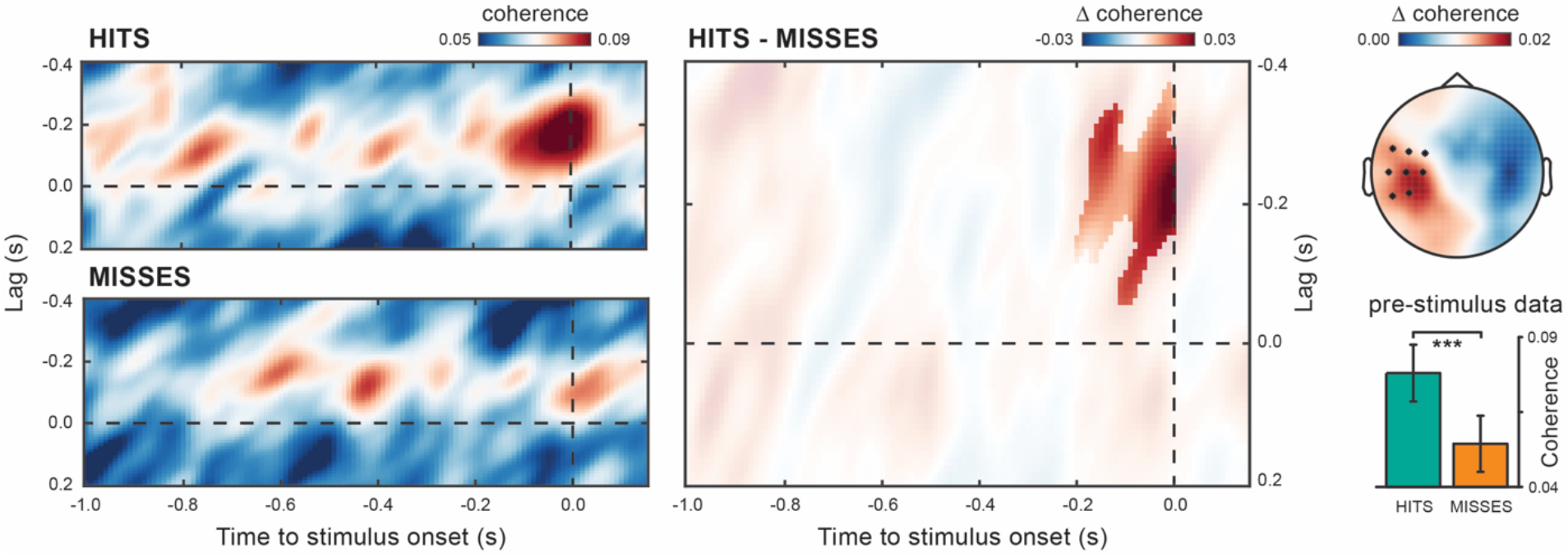
Cortico-force alpha coherence just before stimulus onset predicts perception. Lag- and time-resolved cortico-force alpha (10.5 Hz) coherence over the pre-stimulus period for hits, misses and their difference (hits-misses). The highlighted area indicates the time and lag intervals belonging to the cluster that survived cluster-based permutation statistics for the hits-misses contrast. Alpha coherence is averaged over the EEG electrodes belonging to the same cluster (evaluated at time 0 s and lag −0.2 s; see black asterisks in the topographic map). The topography shows the hits-misses difference in alpha coherence averaged over the time and lag intervals belonging to the same cluster. The bar plot shows alpha coherence for the electrode CP5, calculated at lag −0.2 s and time −0.16 s, i.e., the time point closest to stimulus onset where the analyzed data windows do not include any post-stimulus data point. Error bars indicate ± 1 SEM. \*\*\**p*<0.001.

Cortico-force coupling is strongly enhanced just before the onset time of seen compared to unseen stimuli. This enhancement is selectively observed at negative lags (~-0.2 s) that match well the lag tuning profile of alpha coherence (see above). Cluster-based permutation statistics on pre-stimulus cortico-force coherence confirms a significant difference between hits and misses which is concentrated over left centro-parietal electrodes, at times immediately preceding stimulus onset and for EEG-leading lags (p = 0.0192; corrected for multiple comparisons across time [−1 – 0 s] and lags [−0.4 – 0 s]; cluster time interval: −0.2 – 0 s; lag interval: −0.37 to −0.06 s; Fig 4).

Because of the time-resolved approach, the peri-stimulus coherence estimates are based on (0.3-s) EEG/force data windows which variably (depending on time and lag) embed poststimulus signals. This raises the possibility that the difference in coherence observed close to stimulus onset might reflect detection-related modulations of the responses evoked by seen (hits) and unseen (misses) stimuli (see Fig 1C for the stimulus-evoked responses in the force). To corroborate the ongoing, rather than evoked, nature of the present phenomenon, we therefore restricted the statistical evaluation of coherence to the latest (closest to stimulus onset) time point which encompasses uniquely pre-stimulus data segments (i.e., −0.16 s). As shown in the bar plot in Fig 4, cortico-force coherence evaluated at this (pre-stimulus) time point (and at lag −0.2 s) is significantly stronger for hits compared to misses (cluster p = 0. 0078), excluding any confound due to post-stimulus data contamination.

The perceptual relevance of cortico-force coherence is temporally confined to a short epoch immediately preceding stimulus onset and does not appear as the tail of a longer-lasting modulation. The fast-changing dynamics rules out that the present effect is related to slow fluctuations in arousal or task engagement. Although the (single-trial) temporal dynamics of a phenomenon cannot be directly inferred by looking at its time-locked components, the present results suggest the transient nature of cortico-force coherence and its dynamic influence on perceptual processes.

We also checked that detection-related changes in pre-stimulus oscillatory power could not account for the observed changes in coherence. In line with previous findings (42–44), hits are associated with lower pre-stimulus alpha power compared to misses (the effect does not however reach statistical significance). However, the power and coherence modulations clearly differ both in terms of topography and temporal dynamics (see S2 Fig). Most importantly, the sign of the power modulation (pow _hits_ < pow _misses_) excludes it as a potential confounding factor for the coherence modulation reported herein (coh _hits_> coh _misses_).

### Post-stimulus modulations of cortico-force coherence

We finally checked whether alpha-band cortico-force coherence undergoes stimulus- or detection-related modulations in the post-stimulus window (0-0.5 s).

Fig 5A shows the lag-frequency plot of coherence evaluated at CP5 and averaged over the entire post-stimulus period for all trials combined (left).

**Fig 5.**
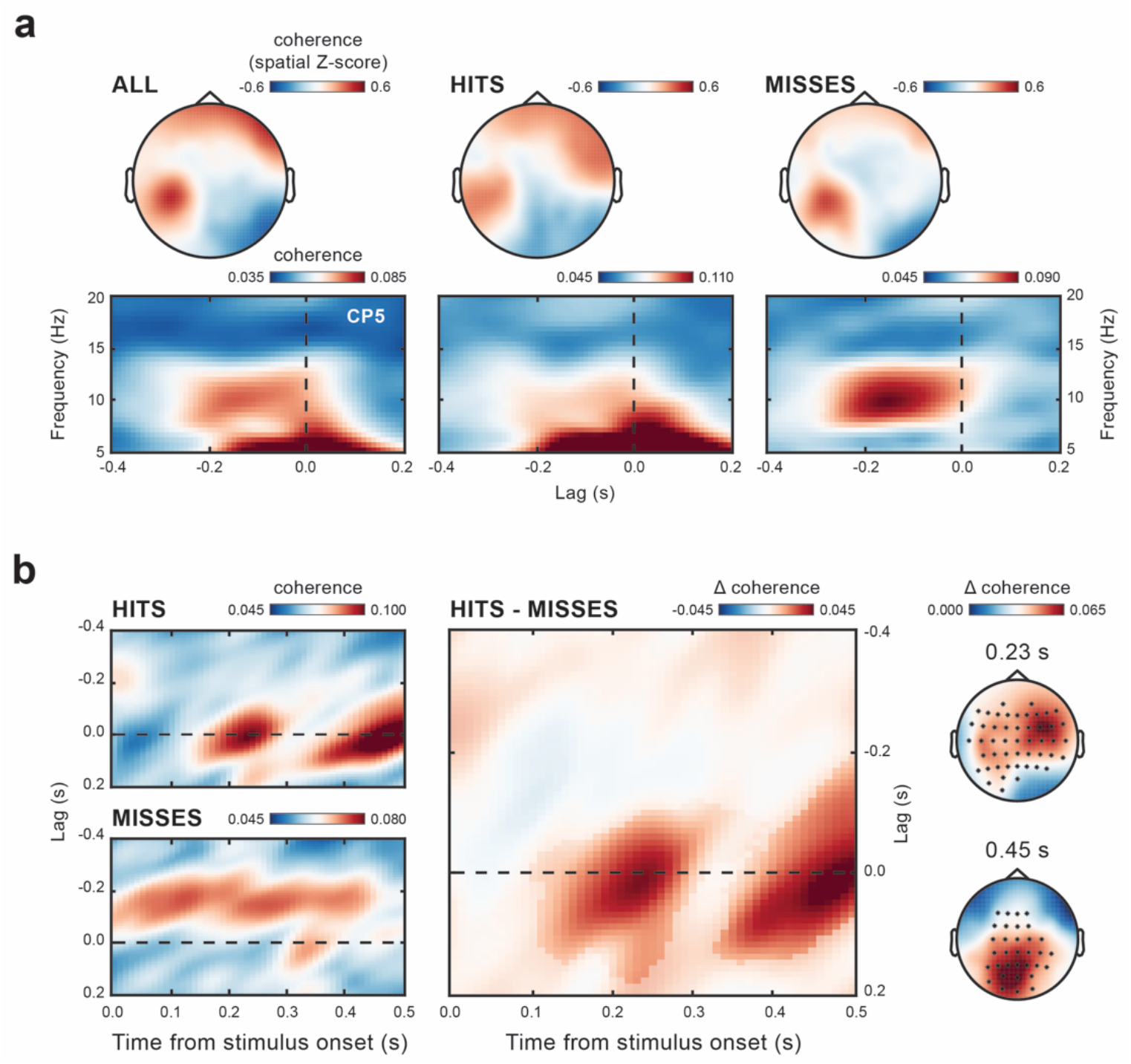
Post-stimulus lagged and zero-lag coherence components. (**A**) Lag-frequency coherence representation as in Fig 3C but averaged over the post-stimulus period (0-0.5 s) for all trials as well as separately for hits- and misses-trials. Topographies show coherence at 10.5 Hz and lag −0.2 s for all trial categories. (**B**) Lag- and time-resolved cortico-force alpha (10.5 Hz) coherence over the post-stimulus period for hits, misses and their difference (hits-misses). The highlighted area indicates the time and lag intervals belonging to two different clusters that survived cluster-based permutation statistics for the hits-misses contrast. Coherence is averaged over all EEG electrodes. Topographies show the hits-misses difference in alpha coherence calculated at zero-lag and at time 0.23 s (where the effect is maximal for the early-latency cluster; top) and 0.45 s (where the effect is maximal for the late-latency cluster; bottom). Black asterisks mark the EEG electrodes belonging to the corresponding clusters.

Like for the pre-stimulus epoch, coherence in the alpha-band increases around negative lags. However, in addition to the alpha lagged component, a marked increase in zero-lag coherence can be observed and is maximal at lower frequencies (<10 Hz). These low-frequency zero-lag components are particularly prominent in the coherence spectrum of hits, spilling over higher frequencies and partly concealing the alpha-selective components (middle). On the contrary, alpha lagged coherence is clearly discernable in the post-stimulus period of misses-trials where the zero-lag components are virtually absent (right). Nonetheless, for both hits and misses alpha coherence evaluated at negative lags shows a topography that closely resembles the one observed in the pre-stimulus window (see Fig 5A and 3E). Thus, alpha coherence appears to conserve nearly identical properties in the pre- and post-stimulus windows, proving as a robust phenomenon.

We also looked at the temporal evolution of alpha coherence after stimulus onset (averaged across channels and subjects; Fig 5B). The hits-related increase in zero-lag coherence is composed of two clusters that differ in terms of latency and topography: an early cluster with ipsilateral frontal distribution (cluster p-value = 0.0036; cluster time interval: from 0.1 to 0.39 s; lag interval: from −0.19 to +0.18 s) and a late cluster with occipital distribution (cluster p-value = 0.011; cluster time interval: from 0.33 to 0.5 s; lag interval: from −0.19 to +0.15 s). Further inspection of their respective spectral properties reveals that the low frequencies only contribute to the late cluster, whereas the early cluster is narrow-band and spectrally confined to the alpha range (see S3 Fig). These zero-lag components likely reflect stimulus-evoked potentials (see also Fig 1C) and/or phase-resetting of rhythmic activity in both the EEG and the force signal. Thus, lag-selectivity allows to discern an ongoing from an evoked alpha component in cortico-force coherence, despite their spectral similarity and temporal coexistence in the post-stimulus period.

Since we are here mainly interested in the ongoing component of alpha coherence that is selectively established at negative lags, we further checked for its possible modulation by statistically contrasting post-stimulus alpha coherence and mean alpha coherence over the −0.5 – 0 s pre-stimulus interval (at electrode CP5 and lag −0.2 s) and found no significant difference at any time point neither for hits nor for misses.

These results corroborate the ongoing nature of alpha coherence, with central alpha leading peripheral alpha in time.

## Discussion

Effective motor behavior entails prediction and continuous monitoring of sensory information. Despite its computational bases have been described by several models (45), little is known about how the coupling between motor and sensory functions is realized at the neural level. The present study shows that the ongoing oscillatory dynamics subtending cortico-motor control is relevant for visual perception, suggesting the operation of a task-independent visuomotor loop.

We report two novel findings. First, 10-Hz fluctuations in the motor output are phase synchronized (coherent) with a cortical alpha rhythm which foreruns the peripheral rhythm in time by about 0.2 s. Second, this non-zero lag motor synchronization predicts visual performance. Stimulus detection is facilitated when cortico-motor alpha synchronization is enhanced immediately before stimulus onset. This suggests that the actual visual gain is upregulated at times when upper and lower sensorimotor centers happen to be in the optimal phase relation for their communication. In accordance with an extensive literature (e.g. 30, 35), we also observed consistent cortico-motor coherence in the beta range (15–30 Hz) which, conversely, peaks around zero lag and bears no relation with visual perception.

### The sensorimotor function of cortico-motor coherence

Coherence between cortical and peripheral (muscle/force) activities – also termed cortico-muscular or corticospinal coherence – is typically greatest in the beta-band and has been largely investigated in relation to motor control. Classically viewed as a somatotopically-organized propagation of oscillatory signals from the motor cortex to its downstream spinal/muscle targets (41, 46), it is now thought to reflect the information flow within a cortico-peripheral-cortical loop, involving both descending and ascending pathways (30). Beta-band coherence with muscle activity has been indeed observed in both human and non-human primates across an extended sensorimotor network, comprising motor (31, 41) and somatosensory (47) cortices, spinal centers (48) and even proprioceptive afferences to the dorsal spinal roots (49). Coherence-based estimates of the time delay between central and peripheral signals have proved extremely variable but more often compatible with bidirectional interactions rather than unidirectional propagation (50). Directed connectivity analyses also support distinct descending and ascending contributions to cortico-motor coupling (48, 51).

Not surprisingly, current accounts of cortico-muscular coherence invariably assign it a major role in the integration of motor and somatosensory signals, being it either for the purpose of keeping the sensorimotor system accurately calibrated (30), maintaining a stable motor output (46, 52, 53), strengthening selective sensorimotor connections, such as those based on muscle synergies (54) or supporting closed-loop sensorimotor control (48). These interpretations collectively discard a passive spillover of oscillatory signals along anatomically connected structures and instead imply a functionally relevant sensorimotor phenomenon. In line with this putative functional role, the strength and spectral properties of cortico-muscular coherence are subject to task-dependent modulations (34) and associated with motor performance (55). For example, beta-band coherence prevails during weak sustained contractions and shifts to higher frequencies (gamma band) during increased force levels (46), dynamic force control (56) and motor preparation (13, 34). However, although a sensorimotor nature of cortico-motor coherence has been postulated (30), its behavioral relevance has only been assessed at the motor, not at the perceptual, level.

### Properties and perceptual relevance of cortico-motor alpha coherence

Overt motor output, i.e., the mechanical result (force/acceleration) of muscle activity, is dominated by lower-frequency fluctuations around 10 Hz (physiological tremor). Evidence that these are related to central neural activity is sparse (37–39, 57) and the origin of physiological tremor is debated (33). Some works have suggested that descending 10-Hz drive may be selectively dampened by phase-cancellation at the spinal level, thereby reducing cortico-motor alpha coupling (58, 59); others have argued for a primary (if not exclusive) contribution of peripheral factors – e.g. mechanical resonance (60, 61) and stretch reflex (62) – over central neural factors in tremor generation.

Here we show that the peripheral alpha rhythm is (at least partly) explained by a corresponding cortical rhythm. Despite the existing debate, the here reported alpha coherence shows spectral, spatial and lag-selective properties which are consistent across different data subsets (hits- and misses-trials; see Fig 3) and support as a whole a genuine central basis of tremor.

An important novelty of the present findings is that continuous 10-Hz force fluctuations are maximally coherent with 0.2-s backward shifted EEG alpha activity, suggesting cortical phase-modulation of the (forthcoming) motor output (a similar phenomenon is observable for cortico-EMG coherence; see supporting information). The standard approaches to study cortico-muscular coherence have either completely disregarded lag-dependence or focused on very short lags (tens of milliseconds) that are inferred from the slope of the phase-frequency function (50). These short lags are also expected from the known conduction times along monosynaptic (or oligosynaptic) corticospinal pathways. This may have contributed to the reports of (close to) zero-lag beta-band coherence instead of lagged alpha-band components. Interestingly, motor cortical spiking activity shows maximal correlation with generated force at an equally long anticipatory lag (i.e., amounting to ~0.2 s) (63). Indeed, when it comes to effective motor control, longer integration times within cortico-subcortical and cerebellar loops might add up to the time needed for an electrical impulse to travel down the corticospinal tract. Thus, alpha lagged coherence uncovers distinct neural processes that are involved in motor control but operate on a slower timescale as compared to those contributing to zero-lag beta coherence.

As voluntary movements are subject to feedback-based control, sensory analysis is inevitably part and parcel of motor processing. Previous works have shown that beta-band local synchronization in motor cortex and muscles (power) as well as along the corticospinal connection (coherence) predict postural holding and speed of movement in response to externally imposed perturbations in proprioceptive (52) and visual (52, 53, 64) feedback. All of these works evaluate sensory-guided motor performance and can therefore hardly disentangle motor and sensory components in neural dynamics and behavior.

The task used in the present study certainly requires the efficient closing of a proprioceptive-motor control loop for its accomplishment. At the same time, motor control is here functionally decoupled from visual processing, as no visual feedback of the exerted force is available. Crucially, we show that the alpha-band oscillatory dynamics in cortico-motor control shapes visual perception. The visual modulation is temporally selective and occurs in the absence of changes in motor performance, ruling out slow fluctuations in arousal or task engagement as major causal factors. Despite the task not demanding it, the current findings reveal that a visuo-motor loop might yet be running in the background and cycling at alpha periodicity with consequences on perception.

### Internal motor dynamics structure sensory sampling

Motor contribution to perceptual processes has been mainly addressed in the context of *active sensing* (65). Indeed, some motor behaviors (e.g. saccadic/haptic exploration) are inherently endowed with a perceptual function. Specifically, when effectors (e.g. the eyes) and sensors (e.g. the retina) share the same anatomical substrate, movements’ net effect is that of relocating sensors in space, de facto supporting overt sensory information sampling.

A recent EEG work looked into the coupling between cortical activity and motor outputs (finger kinematics) during free haptic exploration of textures (66). The same neural activity that was related to the hand movements was also found to be related to the ensuing tactile perception. However, an important difference with our findings is that the spatiotemporal pattern of the incoming (tactile) inputs is unavoidably shaped by the overt (fingers) movements, as it is the case for all exploratory behaviors. Motor exploration (e.g. saccades, whisking, sniffing) often displays a rhythmic pattern. Some authors have proposed that this overt motor dynamics could modulate cortical excitability in a predictable manner, boosting sensitivity for the newly acquired inputs (65, 67). Indeed, monkey data show that visual oscillatory dynamics is entrained to the rhythmic alternation between saccades and fixations (15, 16), and recent human behavioral data support the perceptual relevance of these neural modulations (22, 23, 25).

Most of these perceptual changes that occur at the time of movement have been interpreted as the consequence of adaptive mechanisms aimed at optimizing perception, either by promoting perceptual stability (22) or by enhancing information extraction (27, 65). Surprisingly, a similar phenomenon holds for movements that do not mediate the actual sampling of visual information. It was recently shown that during preparation to move the hand, visual perception undergoes rhythmic fluctuations that are time-locked to the future movement (21). A follow-up experiment revealed that fluctuations in visual perception are explained on a trial-by-trial basis by corresponding brain oscillations phase-locked to action onset (18). Action-perception coupling mechanisms may thus go beyond the presence of a direct causal link between movements and information sampling.

The above set of findings are compatible with the hypothesis that both movement initiation and perceptual sensitivity are jointly regulated by a common, attention-based, rhythmic source (see discussion in 26). Such a rhythmic mechanism tightly fits active exploratory behavior (e.g. eye movements), whereby periods of augmented sensory information extraction would alternate with shifts of attention and movement to new sensory landscapes (27–29). In this framework, sensory and motor processing are coupled (anti-correlated) but yet segregated.

Conversely, the present work seeks to describe the ongoing dynamics of communication between the sensory and motor system which possibly subtends sensorimotor control. We here provide evidence that sensory information sampling is indeed structured by the internal oscillatory motor control dynamics – unambiguously indexed by cortico-force coherence – with observable consequences on perception.

Our finding corroborates the behavioral relevance of neuronal phase synchronization (force can be considered here as a proxy of spinal activity), adding support to its role in effective communication (12). Previous evidence shows that inter-areal (gamma) synchronization within the visual system (V1-V4) predicts the speed of motor responses to visual stimulation (68). Here, we show that visual detection is predicted by synchronization, not in a visual, but in a motor network (the corticospinal circuit) where visual information (possibly) arrives at a late stage as it is transformed into a motor response. Yet, such a visuomotor transformation is not required by the task at hand. This suggests an automatic and covert gating of visual inputs by the ongoing motor control processes. At the physiological level, an interesting possibility is that activity in visual or, at least, perceptually relevant (high-level) areas, is also coupled to some degree to the activity along the corticospinal tract. Alpha cortico-force coherence would thus represent only a partial description of an otherwise more extended functional network. Functional connectivity across the network nodes might also be subject to ongoing fluctuations, as suggested by the transient modulations of alpha synchronization on perception. These are fascinating hypotheses that can be experimentally addressed in future work. In the following, we would like to speculate about the possible functional significance of our findings in relation to sensorimotor control.

Motor behavior commonly relies on many sources of information which must be integrated in time and space to achieve an efficient control. Beta- and alpha-band cortico-motor coherence may represent the neural signature of multimodal sensorimotor loops operating in parallel at different timescales and to different purposes. Sensorimotor control depends on quickly updated state estimates of the effectors. In line with previous proposals (30, 48), (zero-lag) beta coherence could serve the operation of a proprioceptive-motor loop through which information is fed forward and back at a relatively fast rate.

Actions, in real-world scenarios, require monitoring of task-relevant sensory feedback but also being ready to adapt the motor plan to new data coming from all the senses. Incorporating visual, and possibly multimodal, information into the ongoing motor plans would require however longer integration times. Even when not task-relevant, visual information could be (covertly) monitored within a slower control loop, possibly indexed by 0.2-s lagged alpha coherence. Of course, this does not exclude the possibility that the strength and/or properties (e.g. lag-tuning) of alpha coherence are modulated in a context-dependent fashion (e.g. by the importance of visual feedback for motor control).

In conclusion, we suggest that the online monitoring of multimodal sensory information is integral to the organization and control of movement and may be indexed by different sensorimotor communication channels (alpha vs. beta coherence) running with different lags. Such a lag specificity may stem from the inherent functional role played by proprioceptive vs. exteroceptive signals in tuning the descending command. The former allows a fine and short latency control of muscle activation via direct modulation of spinal reflexive circuitries. The latter, by requiring a transcortical path, is endowed with far less potential to control the details and timing of muscle recruitment in favor of a more general role in guiding actions at higher levels of the cognitive hierarchy.

Oscillatory mechanisms could thus serve to synchronize incoming inputs with descending motor commands/predictions, effectively regulating the information flow within multi-modal and multi-timescale sensorimotor loops.

## Materials and Methods

### Subjects

Twenty-five healthy participants were recruited to participate in the experiment. Due to excessive difficulty in the performance of the task (specifically in isometric force control; see below), five participants did not complete the experiment but only attended one out of the three testing days; the data for these participants were not analyzed.

The remaining twenty participants (11 females; age 22.1 ± 3.2 years, MEAN±SD) took part to the full experiment. Participants were all naïve with respect to the aims of the study and were all paid (€12.5/testing day) for their participation. Participants were right-handed (by selfreport) and had normal or corrected-to-normal vision. The study and experimental procedures were approved by the local ethics committee (Comitato Etico della Provincia di Ferrara, approval number: 170592). Participants provided written, informed consent after explanation of the task and experimental procedures, in accordance with the guidelines of the local ethics committee and the Declaration of Helsinki.

### Experimental setup and procedure

Participants sat in a dimly lit room in front of a CRT screen (21”, 85 Hz; Sony Trinitron Multiscan-500PS) at a viewing distance of ~60 cm. They held a dual-axis custom-made isometric joystick with their right hand that allowed measuring hand force continuously along two orthogonal axes via four load cells. The joystick was securely fixed to a rigid support to avoid displacement and enclosed in a black casing with an aperture in the front to prevent participants from seeing their hand.

Participants performed concurrently a motor task and a visual detection task. The continuous isometric motor task consisted in a wrist abduction to push the joystick’s handle towards one’s own body with the right hand. The force level that participants were required to exert (i.e., target force) was set at the beginning of the experiment based on the individual maximal voluntary force (MVF; see below).

Each trial was structured as follows. A dark grey fixation cross (size 0.35°) was first displayed at the center of the screen. Participants were instructed to start exerting force as soon as they felt ready. As participants began pushing the joystick, four horizontal bars (left/right and top/bottom of fixation; horizontal/vertical eccentricity: 7.5°; see Fig 1B) linearly increased their length towards the center of the screen as a function of the applied force, providing on-line visual feedback of the exerted force. The target force was reached when the left and right bars (top and bottom bars) met at the center of the screen (see Fig 1B). As soon as the participant succeeded in maintaining force within the desired range (target force ± 0.15*target force) for at least 1.5 s, the bars stopped changing length and remained precisely aligned to the center of the screen. From this moment onwards (i.e., when the visual feedback of the force was removed), participants were instructed to keep contraction as constant as possible without feedback for 5.5 s and, at the same time, pay attention to the appearance of a brief (0.012 s; one frame) visual dot with near-threshold contrast. The dot (size 7’ of visual angle) was shown 7.5° to the right of fixation in 85% of the trials; in the remaining 15% of the trials (catch trials), no visual stimulus was displayed. Importantly, the stimulus was unpredictable in time, as it appeared at a time that was randomly drawn from a uniform distribution ranging between 1.6 and 4.6 s (i.e., 3 s jitter) after fulfillment of the force criterion and consequent visual feedback removal. At the end of the trial (after 5.5 s), indicated by a question mark at the center of the screen, participants released hand contraction and reported verbally whether they had seen (‘yes’ response) or not seen (‘no’ response) the stimulus.

Prior to the experiment, we estimated first the individual MVF and then the individual visual contrast threshold, i.e., the contrast yielding 50% of ‘yes’ responses (for stimulus-present trials).

### Maximal Voluntary Force (MVF) estimation

Participants were asked to apply their maximal force by pushing the joystick handle backward with their right hand in response to a beep (800 Hz, 0.05 s) and maintain the same force for 3 s (end of interval marked by a second identical beep). This procedure was repeated three times with a 7-s pause in-between repetitions. MVF was estimated as the mean force during the 3-s interval averaged over at least two repetitions in which mean force did not differ (across repetitions) by more than 5%. The entire procedure was repeated until this criterion was satisfied.

The lower is the force to be exerted the more difficult is its fine control and stabilization; at the same time, high forces induce fatigue over the course of the experiment. For these reasons, the target force used for the experiment was set as the minimum force between 10 and 20% of MVF at which the participant was capable of controlling the joystick with no excessive difficulty. Except from five subjects who withdrew before completing the experiment, no other subject reported discomfort or fatigue throughout the experiment.

### Visual contrast threshold estimation

Task and trial structure were the same as already described for the main experiment. The contrast of the visual stimulus was changed on a trial-by-trial basis according to the adaptive QUEST algorithm (69). We run 60 trials and fitted the obtained data with a cumulative Gaussian function. The contrast threshold was estimated as the mean of the psychometric function.

Due to possible learning effects, the performance level (hit rate, i.e., percentage of ‘yes’ responses for stimulus-present trials) in the main experiment was continuously monitored and the stimulus contrast was adjusted throughout the experiment to keep performance near threshold. The hit rate was calculated at the end of each block of trials (i.e., 60 trials). The contrast used in the next block of trials was not changed if hit rate was between 45 and 55%. The contrast was decreased/increased by 0.4 dB if hit rate was within 55–65% or 35–45%, respectively, by 0.8 dB if it was within 65–75% or 25–35%, and by 1.2 dB if it was >75% or <25%.

A photodiode (2.5 x 2.5 cm) was placed in the bottom right corner of the monitor and was used to align the data with millisecond accuracy. Specifically, a white square was displayed on the screen at the position of the photodiode (hidden from view) in synchrony with the removal of the force-related visual feedback as well as at the end of the trial.

Both the voltage signal from the photodiode and that from the isometric joystick were recorded by a data acquisition board (USB-1608GX, Measurement Computing; sampling rate, 5000 Hz). The presentation of the stimuli and the data acquisition device were controlled via Matlab Psychtoolbox-3 (RRID: SCR_002881).

### Data collection

Data collection was split in three different testing days (2 hrs. testing each day). The experiment involved separate blocks of 60 trials each, with few minutes of rest in-between blocks. Participants completed on average a total of 654±53 (MEAN±SD) trials.

### EEG and EMG recording

EEG data were recorded continuously during the experiment (except during MVF and visual contrast threshold estimation) with a 64-channel active electrode system (Brain Products GmbH, Gilching, Germany). Electrooculograms (EOGs) were recorded using four electrodes from the cap: FT9, FT10, PO9 and PO10 were removed from their original scalp sites and placed at the bilateral outer canthi and below and above the right eye to record horizontal and vertical eye movements, respectively.

All electrodes were on-line referenced to the left mastoid. The impedance of the electrodes was kept below 15 kΩ. EEG signals were sampled at 1000 Hz.

Additionally, EMG was collected from a right arm muscle recruited in the radial deviation of the wrist (Extensor Carpi Radialis Longus – ECRL), a synergic muscle for wrist abduction (Flexor Carpi Radialis – FCR) and a control muscle (Biceps Brachii Brevis – BBB). Muscles of interest were located via standard palpation procedures and montage was on the muscle belly with a ~3-cm interelectrode distance. EMG was recorded with a wireless system (Zerowire EMG, Aurion, Italy), acquired by a CED board (Micro1401; sampling rate, 5000 Hz) and visualized online with the Signal 3.09 software (Cambridge Electronic Design, Cambridge, UK).

The signal from the photodiode was converted in a TTL signal by an Arduino Due board and used to accurately synchronize all acquired data (EEG, EMG, Force) and relevant task events.

### Data analysis

Analyses of behavioral and EEG/EMG data were performed with custom-made Matlab code and the FieldTrip toolbox [(70); RRID:SCR_004849].

Data were first temporally aligned to the visual stimulus presentation and then epoched in 2.5 s-long segments from −1.6 to 0.9 s (relative to stimulus onset).

Force traces were visually inspected, and trials were rejected if the force was at any time (during the relevant epoch) less than 20% of the individual target force.

EEG/EMG segmented data were manually checked for bad channels and/or artifacts in the time domain. Trials containing eye movements (saccades, blinks) within a 0.25-s window before stimulus onset were discarded from the analysis. Independent Component Analysis (ICA) was used to identify and remove residual artifacts in the EEG signal related to eye movements and heartbeat. Bad EEG channels were excluded from the ICA analysis and subsequently interpolated with a distance-weighted nearest-neighbor approach. Fp1, Fp2, AF7 and AF8 were excluded from the analysis due to greater noise level in a large number of subjects.

Both EMG and Force were down sampled to 1000 Hz.

### Behavioral analysis

Behavioral performance was evaluated on the force, because it was the output parameter that the subject was requested to control throughout the task. Force was first low-pass filtered (35-Hz cutoff frequency) and then analyzed separately in the pre- (−1.6 – 0 s) and post-stimulus (0 − 0.85 s) windows.

Pre-stimulus force values were expressed as the percentage change relative to the individual target force ([force – target force]/[target force]*100); these values and their modulus were averaged over time (−1.6 – 0 s) to yield an estimate of motor performance accuracy, i.e., force deviation and absolute error, respectively. Within- and inter-trial force variability were calculated as the standard deviation across time and across trials, respectively. Linear trends were analyzed by submitting (single-trial) pre-stimulus force to a linear least-squares fitting and deriving the slope of the best-fitting functions (Fig 1D).

For the analysis on the post-stimulus window, we first detrended the force based on the −0.5 – 0 s pre-stimulus interval and then applied baseline correction using the −0.05 to 0 s pre-stimulus interval (in accordance with (32); Fig 1C).

### Spectral analysis

The frequency content of Force and its coherence with cortical (EEG) activity was first analyzed on the entire pre-stimulus window (−1.6 – 0 s) for frequencies between 5 and 35 Hz (0.5-Hz step) by using Fourier-based analysis combined with the multi-taper method [(71); 3- Hz spectral smoothing; Fig 2].

Lagged coherence was computed by applying short-time Fourier transform on hanning tapered data segments. In a first analysis, we used a 0.6-s data window (from −1.1 to −0.5 s) and computed coherence by systematically shifting the EEG signal (relative to the Force signal) by different amounts of time: from −0.5 s (EEG precedes Force by 0.5 s; i.e., EEG data window: – 1. 6 – −1 s) up to +0.5 s (EEG follows Force by 0.5 s; i.e., EEG data window: −0.6 – 0 s) in 10- ms steps (Fig 3A).

Next, we computed lagged coherence with a time-resolved approach using a 0.3-s sliding window that was advanced over the data from −1 to −0.35 s (relative to stimulus onset) by 10- ms steps. To zoom into the phenomenon of interest (i.e., alpha-band cortico-force coherence), this analysis was restricted to frequencies between 5 and 20 Hz and lags from −0.4 to 0.2 s (Fig 3C,D,E).

Finally, we contrasted time-resolved lagged coherence for hits- and misses-trials. Based on the previous analyses, we selected the frequency showing maximal coherence over CP5 for all trials. Coherence was then computed for the previously selected frequency (i.e., 10.5 Hz) and for lags from −0.4 to 0.2 s using a 3-cycles (i.e., ~0.286 s) sliding window advanced over the data from −1 to 0.15 s (Fig 4).

Equivalent analyses have been produced for the post-stimulus window (from 0 to 0.5 s; see Fig 5A,B).

Cross-correlation of EEG and Force (normalized so that the autocorrelations at zero lag are identically 1) was computed on 1-s data segments (from −1 to 0 s) that were previously bandpass filtered in the beta (20–30 Hz) and alpha (8–12 Hz) range with a two-pass Butterworth filter (2^nd^ order for each single pass; Fig 3B).

### Statistical analysis

Motor performance (accuracy and variability) was compared between hits- and misses-trials using conventional paired samples t-tests.

All other statistical comparisons between hits- and misses-trials were performed using cluster-based permutation tests (72) that allow to deal more effectively with the multiple comparisons along the spatial, frequency and time dimensions. According to this non-parametric statistical approach, all samples exceeding an a priori decided threshold (uncorrected p < 0.05, two-tailed) for univariate statistical testing (dependent-sample t-test) are selected and subsequently clustered on the basis of their contiguity along the relevant (spatial, spectral and temporal) dimension(s). Cluster-level statistics is computed by taking the sum of t-values in each cluster. This sum is then used as test statistic and evaluated against a surrogate distribution of maximum cluster t-values obtained after permuting data across conditions (at the level of participant specific condition averages). To generate the surrogate distribution, we used 5000 permutations. The p-value is given by the proportion of random permutations that yields a larger test statistic compared to that computed for the original data.

## Supporting Information captions

**S1 Fig. Cortico-EMG coherence.**

Lag- and frequency-resolved coherence between EEG and EMG activity recorded from a wrist extensor muscle (Extensor Carpi Radialis Longus – ECRL). Lagged cortico-EMG coherence is calculated on 0.6-s data windows in the same way as shown in Fig 3A for cortico-force coherence. The raw EMG has been high-pass-filtered at 10 Hz and rectified before computing coherence. Cortico-EMG coherence shows similar properties as those observed for cortico-force coherence. Coherence in the beta-band (20–30 Hz) is symmetrically distributed around zero lag and is maximal over contralateral central electrodes (left topography; C1 marked in grey). Coherence in the alpha-band (9–11 Hz) is biased towards negative lags and distributed over contralateral centro-parietal electrodes (right topography; CP5 marked in grey). Notably, alpha-band cortico-EMG coherence peaks at a slightly shorter lag (~−0.14 s) compared to its cortico-force counterpart (~-0.2 s) which is compatible with the fact that muscular activity anticipates in time its mechanical outcome (i.e., force).

**S2 Fig. Pre-stimulus modulations in EEG alpha power do not account for cortico-force coherence modulations.**

Time-frequency plot of power modulations (hits/misses) averaged over selected occipital EEG electrodes (marked in grey in the topographic map). The topography shows the hits-misses difference in power calculated at stimulus onset (0 s) and at frequency 10.5 Hz.

**S3 Fig. Post-stimulus modulations in zero-lag cortico-force coherence.**

Time-frequency plot of post-stimulus coherence at lag 0 s, averaged over all EEG electrodes and shown separately for hits (top) and misses (bottom).

